# Effect of shipping on the microbiome of donor mice used to reconstitute germ-free recipients

**DOI:** 10.1101/2024.03.29.587359

**Authors:** Zachary L. McAdams, Jared Yates, Giedre Turner, Rebecca A. Dorfmeyer, Mary Wight-Carter, James Amos-Landgraf, Craig L. Franklin, Aaron C. Ericsson

**Affiliations:** Mutant Mouse Resource and Research Center at the University of Missouri (MU MMRRC), Columbia, MO 65201; MU Metagenomics Center (MUMC), Columbia, MO 65201; School of Medicine, University of Missouri (MU), Columbia, MO, 65212; UT Southwestern Medical Center, Dallas, TX 75390; Department of Veterinary Pathobiology, MU College of Veterinary Medicine, Columbia, MO 65201

**Keywords:** Gut microbiome, 16S rRNA, Jackson Laboratory, Envigo Laboratory, germ-free, fecal microbiota transfer, FMT

## Abstract

The gut microbiota (GM) influences multiple processes during host development and maintenance. To study these events, fecal microbiota transfer (FMT) to germ-free (GF) recipients is often performed. Mouse models of disease are also susceptible to GM-dependent effects, and cryo-repositories often store feces from donated mouse strains. Shipping live mice may affect the GM and result in an inaccurate representation of the baseline GM. We hypothesize that the use of such fecal samples for FMT would transfer shipping-induced changes in the donor GM to GF recipients. To test this, donor mice originating from two suppliers were shipped to the University of Missouri. Fecal samples collected pre- and post-shipping were used to inoculate GF mice. Pre- and post-shipping fecal samples from donors, and fecal and/or cecal contents were collected from recipients at one and two weeks post-FMT. 16S rRNA sequencing revealed supplier-dependent effects of shipping on the donor microbiome. FMT efficiency was independent of shipping timepoint or supplier, resulting in transmission of shipping-induced changes to recipient mice, however the effect of supplier-origin microbiome remained evident. While shipping may cause subtle changes in fecal samples collected for FMT, such effects are inconsistent among supplier-origin GMs and minor in comparison to other biological variables.

## Introduction

The gut microbiome (GM) affects numerous host developmental processes, and is essential for long-term host health. While a wide range of different community compositions can perform comparable function during health^1^, sufficient change (i.e., dysbiosis) is often associated with disease^2^. Mice used in biomedical research are traditionally colonized with a specific pathogen-free (SPF) microbiome, and mice purchased from different suppliers harbor distinct supplier-origin (SO) GMs^3–6^. Following purchase from one of the suppliers of SPF mice, myriad husbandry and institution-specific factors^7–9^, as well as shipping itself^10^, can influence the composition of the microbiome at an individual and colony level. As in human health, the phenotype of many mouse models are influenced by specific features within the GM. Indeed, there are numerous reports of changes or even complete loss of phenotype in mouse models in association with a change in the microbiome due to institution-specific practices^11–14^.

Periodic collection and preservation of fecal samples from a research colony allows for benchmarks against which samples can be compared in the future, in the event of changes in model phenotypes. Ostensibly, such samples could also be used to re-inoculate mice via fecal microbiota transfer (FMT) in efforts to restore microbiome-dependent phenotypes. Related to this, the MU NIH-funded Mutant Mouse Resource & Research Center (MMRRC) at the University of Missouri (MU) and other consortium members routinely collect feces from mouse strains donated for cryopreservation, in an effort to preserve the GM at the time of strain curation. While some donating investigators collect and submit fecal samples alongside mice submitted for cryopreservation, the MU MMRRC collects samples from donated mouse strains after arrival at our facility and prior to cryopreservation. Considering the potential effect of shipping mice for cryopreservation on the composition of fecal samples collected for posterity, we wanted to investigate whether shipping mice to be used as donors significantly affects the composition of their GM, and if so, whether any shipping-induced changes in the GM of donors are maintained following reconstitution of germ-free mice.

To do so, fecal samples were collected from donor mice pre- and post-shipping, and used to reconstitute recipient mice via FMT. FMT can be performed using germ-free (GF) or antibiotic-treated recipient mice. To eliminate potential effects of variability among antibiotic-treated recipient mice, GF recipients were used. Lastly, to determine whether any observed effects of shipping are consistent across different SPF microbiomes, donor mice from two suppliers with distinct SO GMs were used. Fecal samples collected from C57BL6/J and C57BL/6NHsd donors pre- and post-shipping, and from recipient GF Swiss Webster mice at one and two weeks post-FMT, were subjected to DNA extraction, 16S rRNA library preparation and sequencing, and testing for the effects of shipping on the GM of mice to be used as FMT donors.

## Methods

### Mice

Donor mice used in the current study were either female, adult (5-6 week-old) C57BL/6J (Jackson Laboratory, *n* = 12) or C57BL/6NHsd (Envigo, *n* = 12) mice. Prior to the study (during the acclimation period), mice were housed in individually ventilated microisolator cages (Allentown) on a 14:10-h light:dark cycle. Recipient mice were germ-free (GF), female, adult (7-8 week-old) Swiss Webster mice (Taconic, *n* = 24), shipped from the supplier in microisolator cages within a flexible film isolator to maintain GF status. Following inoculation of GF recipient mice via FMT, mice were housed in individually ventilated microisolator cages (Thoren, Hazleton, PA) on a 14:10-h light:dark cycle. Temperature was maintained at 22 ± 2°C, with a relative humidity of 30% to 70%. Mice were individually housed on PAPERCHIP® bedding (Watertown, TN) with *ad libitum* access to commercial rodent diet (Formulab Diet 5008, Purina) and autoclaved, acidified water. Autoclaved nestlets were supplied for each cage for additional enrichment. All sentinel mice monitoring these colonies were consistently seronegative for MHV, MVM, MPV, Parvovirus NS-1, TMEV, Murine rotavirus, *Mycoplasma pulmonis*, and Sendai virus. PCR testing was negative for fur mites and pinworms.

### Experimental design

Donor mice were shipped from their respective suppliers directly to UT Southwestern Medical Center (UTSMC). Mice were allowed to acclimate for two weeks, to mitigate any effects of that initial shipment on the GM. Following that, fecal samples were collected from all donor mice the morning of the day they were shipped overnight from UTSMC to the University of Missouri (MU). Fecal samples collected pre-shipping were also shipped overnight to MU on dry ice. Upon arrival at MU, fecal samples were again collected from all donor mice as they were unpacked. To control for sample preservation, these post-shipping fecal samples were also frozen immediately after collection and kept at -80°C until ready for use.

GF recipient mice were shipped directly to MU. Upon arrival, mice were randomly assigned (using a random number generator) to the pre-or post-shipping group. As they were unpacked from the flexible film isolator, mice then received their assigned FMT using donor material collected pre- or post-shipping from UTSMC to MU. Recipient mice were then individually housed for two weeks, with fecal sample collection occurring at 7 and 14 days post-FMT.

### Fecal microbiota transfer

Working in a biosafety cabinet, fecal pellets were removed from the freezer. To prepare donor fecal material for fecal microbiota transfer (FMT), one fecal pellet per procedure was placed in a 2 mL round-bottom tube containing a 0.5 cm diameter stainless steel bead and 500 µL of sterile phosphate-buffered saline (PBS) and agitated at 1/30s using a TissueLyser II (Qiagen) for 60 seconds. Following a pulse centrifugation, the supernatant was carefully collected using a sterile 1 mL pipette and transferred to a nylon mesh filter with a 40 µm pore size and allowed to flow into a collection tube. Collection tubes containing filtered FMT material were then sealed and promptly transported to animal rooms for FMT procedures.

### DNA extraction

Fecal DNA was extracted using PowerFecal kits (Qiagen) according to the manufacturer instructions, except that samples were homogenized in bead tubes using a TissueLyser II (Qiagen) for ten minutes at 30/sec, instead of using the vortex adapter described in the protocol, before proceeding according to the protocol and eluting in 100 µL of elution buffer (Qiagen). DNA yields were quantified via fluorometry (Qubit 2.0, Invitrogen, Carlsbad, CA) using quant-iT BR dsDNA reagent kits (Invitrogen) and normalized to a uniform concentration and volume.

### 16S rRNA library preparation and sequencing

Library preparation and sequencing were performed at the University of Missouri (MU) Genomics Technology Core. Bacterial 16S rRNA amplicons were constructed via amplification of the V4 region of the 16S rRNA gene with universal primers (U515F/806R) previously developed against the V4 region, flanked by Illumina standard adapter sequences^15,16^. Dual-indexed forward and reverse primers were used in all reactions. PCR was performed in 50 µL reactions containing 100 ng metagenomic DNA, primers (0.2 µM each), dNTPs (200 µM each), and Phusion high-fidelity DNA polymerase (1U, Thermo Fisher). Amplification parameters were 98°C^(3 min)^ + [98°C^(15 sec)^ + 50°C^(30 sec)^ + 72°C^(30 sec)^] × 25 cycles + 72°C^(7 min)^. Amplicon pools (5 µL/reaction) were combined, thoroughly mixed, and then purified by addition of Axygen Axyprep MagPCR clean-up beads to an equal volume of 50 µL of amplicons and incubated for 15 minutes at room temperature. Products were then washed multiple times with 80% ethanol and the dried pellet was resuspended in 32.5 µL EB buffer (Qiagen), incubated for two minutes at room temperature, and then placed on the magnetic stand for five minutes. The final amplicon pool was evaluated using the Advanced Analytical Fragment Analyzer automated electrophoresis system, quantified using quant-iT HS dsDNA reagent kits, and diluted according to Illumina’s standard protocol for sequencing as 2×250 bp paired-end reads on the MiSeq instrument.

## Informatics analysis

Sequences were processed using the Quantitative Insights into Molecular Ecology 2 v2021.8^17^ framework. Paired-end reads were trimmed of the universal primers and Illumina adapters using *cutadapt*^18^. Reads were then denoised into unique amplicon sequence variants (ASVs) using DADA2^19^ with the following parameters: 1) reads were truncated to 150 bp in length, 2) reads with greater than 2 expected errors were discarded, 3) reads were merged with minimum overlap of 12 bp, 4) chimeras were removed using the ‘consensus’ method. Unique sequences were filtered to between 249 and 257 bp in length. The resulting feature table was rarefied to an even depth of 43,675 features per sample. PAST^20^ v.4.03 was then used to evaluate alpha (within-sample) and beta (between-sample) diversity.

## Statistical Analysis

Differences in univariate data (i.e., Chao1 and Shannon indices) were assessed using two-way analysis of variance (ANOVA). Tukey *post hoc* testing was performed to identify differences between individual groups. Differences in multivariate data (i.e., beta diversity) was assessed using permutational multivariate analysis of variance (PERMANOVA). Differences in beta diversity were visualized using a principal coordinate analysis (PCoA) of a quarter-root transformed feature table. Beta diversity was compared using Bray-Curtis (weighted) and Jaccard (unweighted) distances.

## Results

### Shipping subtly affects the donor microbiome but supplier-origin differences remain dominant

Comparison of the richness of C57BL/6J (B6J) and C57BL/6NHsd (B6N) donors pre- and post-shipping revealed the expected supplier-origin effect (*p* < 0.001, F = 411) and a significant, albeit much smaller, effect of shipping (*p* = 0.012, F = 6.8) with no interaction between effects (**Figure 1a**). Pairwise comparisons failed to detect significant time-dependent differences within either substrain independently. There were similarly weighted effects of supplier (*p* < 0.001, F = 874.5) and shipping (*p* = 0.019, F = 5.9) on Shannon diversity, and no interaction (**Figure 1b**).

**Figure 1.**
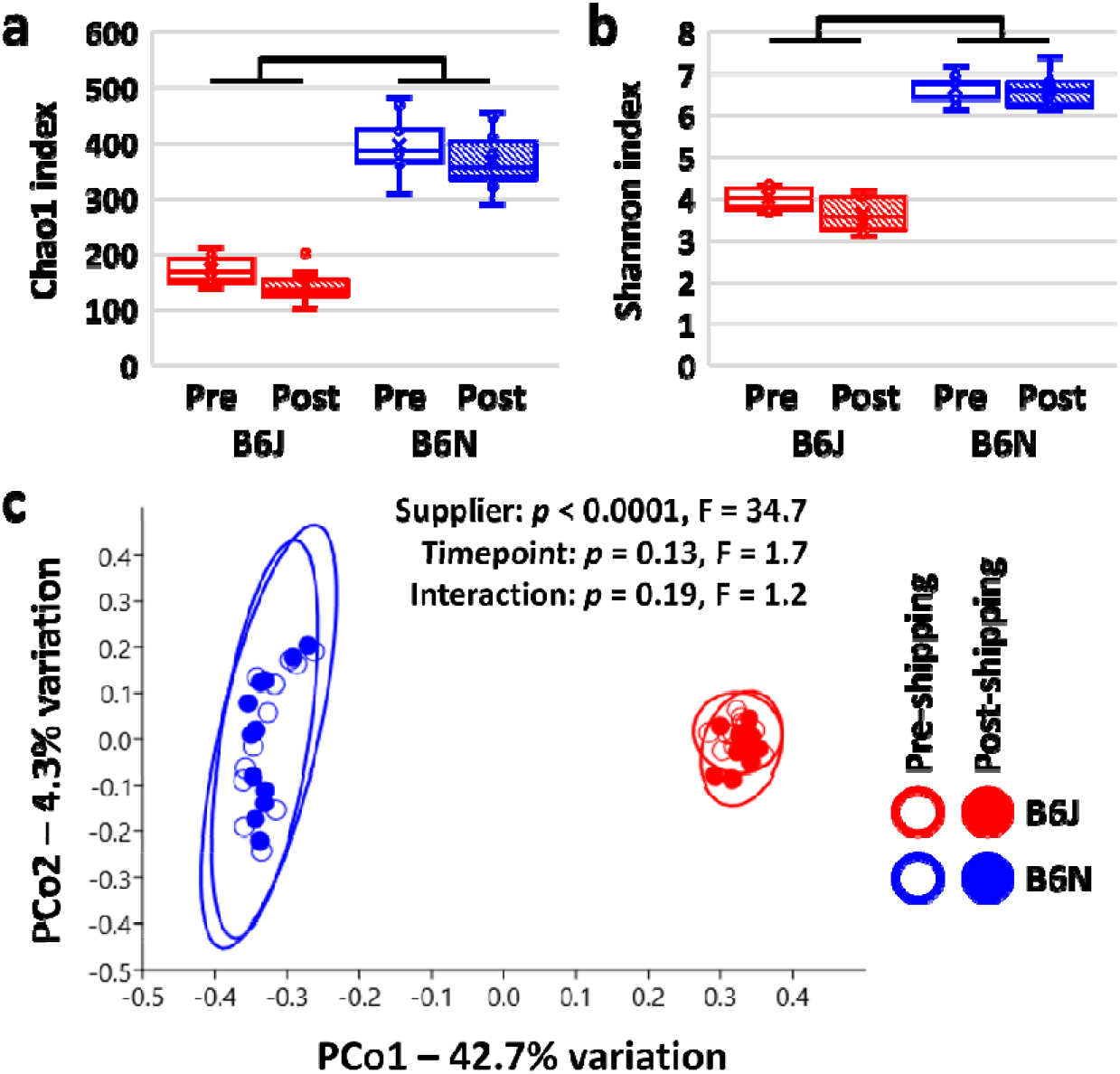
Box plots showing the pre- and post-shipping Chao1 index **(a)** and Shannon diversity index **(b)** in C57BL/6J (B6J) and C57BL/6N (B6N) donor mice originally purchased from Jackson Laboratory and Envigo respectively; Bars denote p < 0.001, two-way ANOVA. Principal coordinate analysis plot **(c)** based on Jaccard distances, showing separation of B6J and B6N donors, and overlap within each substrain of samples collected pre- and post-shipping; legend at right. Ellipses indicate 95% confidence intervals. Results of two-way PERMANOVA using Jaccard distances are shown.

Comparison of beta-diversity via principal coordinate analysis (PCoA) using Jaccard distances revealed clear separation of supplier-origin (SO) GMs, but negligible separation of pre- and post-shipping samples (**Figure 1c**). Two-way PERMANOVA revealed a significant effect of SO GM (*p* < 0.0001, F = 34.7) and no significant effect of shipping (*p* = 0.13, F = 1.7) or interaction. Using Bray-Curtis distances, PCoA showed a similar pattern but PERMANOVA now detected a subtle effect of shipping on the donor microbiome (*p* = 0.04, F = 3.8) and interaction between supplier and timepoint (*p* = 0.04, F = 3.6, **Supplemental Figure S1)**.

### Reconstitution of germ-free recipients results in reduced richness but equivalent alpha-diversity

Following inoculation of germ-free (GF) recipients with donor fecal material collected pre- or post-shipping, recipient mice were maintained under barrier conditions and fecal samples were collected at one and two weeks post-FMT. Cecal contents were also collected at two weeks post-FMT. In GF mice receiving FMT from pre- and post-shipping B6J mice, microbial richness reached between 50 and 60% of donor richness by one week post-FMT and failed to increase beyond that in feces or cecum two weeks later (**Figure 2a**). One recipient of FMT with material from a post-shipping B6J failed to develop richness on par with others in the group, and two-way ANOVA detected significant effects of shipping (*p* = 0.009, F = 7.8) and donor vs recipient (*p* < 0.001, F = 22.1), with no interaction. Pairwise post hoc comparisons found significantly lower richness in recipients at all timepoints compared to their respective donor group. In contrast, the richness of mice receiving FMT from pre- and post-shipping B6N donors reached 50-60% of donor richness by one week post-FMT but continued to develop before reaching 70-80% of donor richness by two weeks, despite the B6N microbiome being over twice as rich as the B6J microbiome. Here, two-way ANOVA failed to detect an effect of shipping (*p* = 0.965, F = 0) but did detect an overall difference between donors and recipients (*p* < 0.001, F = 24.1), and no interaction (**Figure 2b**). Pairwise comparisons found reduced richness in recipients of pre-shipping material relative to donors at all timepoints. In recipients of post-shipping B6N material, fecal richness failed to achieve donor richness, but cecal contents of recipients were not different from donor material.

**Figure 2.**
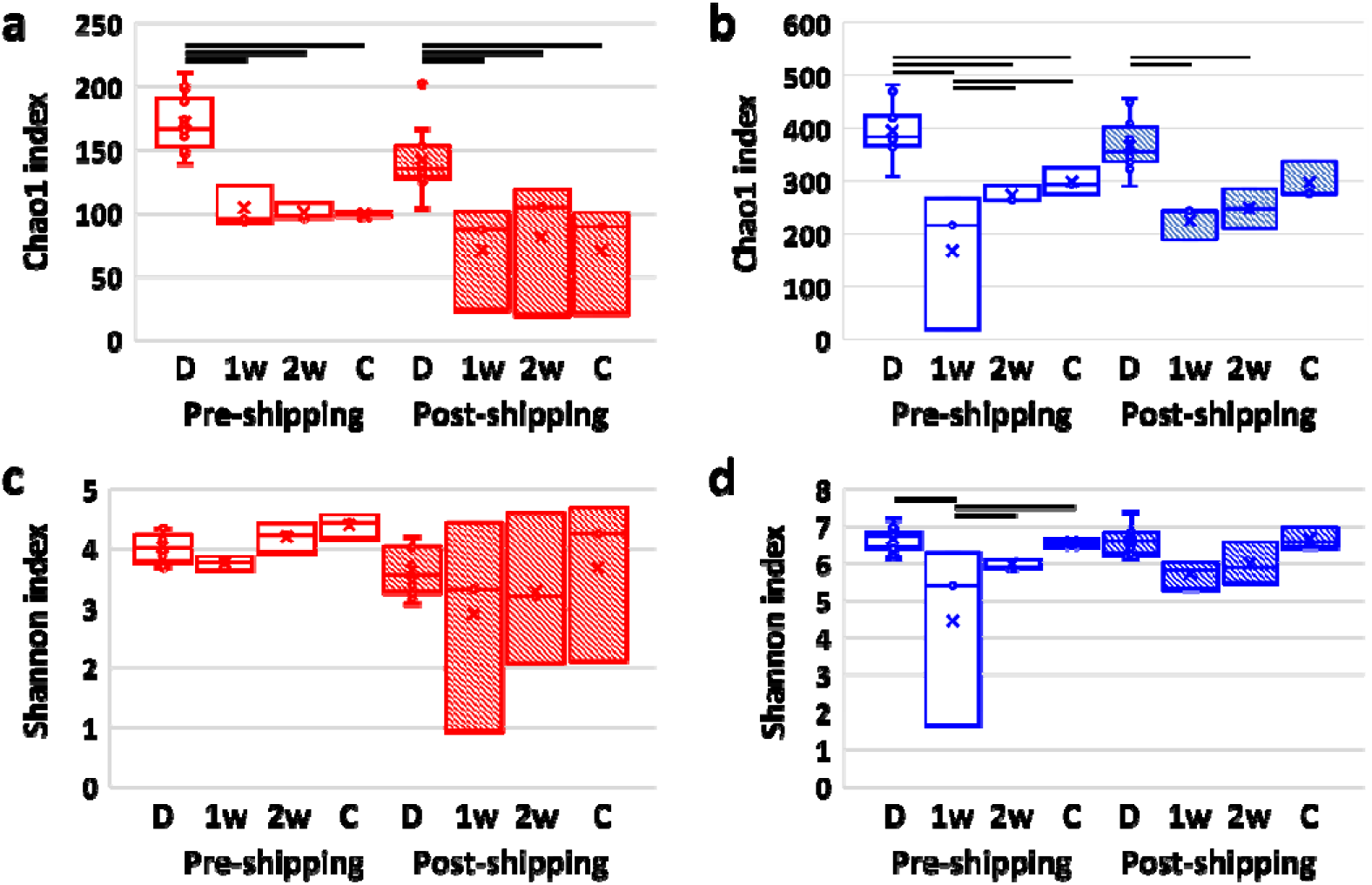
Box plots showing the Chao1 index in feces of ex-germ-free fecal microbiome transfer (FMT) recipients of material from B6J donors **(a)** or B6N donors **(b)** one week (1w) or two weeks (2w) post-FMT, or cecal contents (C) two weeks post-FMT, in relation to the relevant donor (D) group. **(c, d**) Box plots showing Shannon diversity in the same groups. Bars represent significantly differing groups. *p* < 0.05 Tukey *post hoc* testing.

Similar comparison of the Shannon diversity revealed a significant effect of shipping on B6J donors and their recipients (*p* = 0.007, F = 8.3) but no overall (*p* = 0.322, F = 1.2) or pairwise differences between donors and recipients and no interactions (**Figure 2c**). In mice receiving B6N material however, there was no effect of shipping on Shannon diversity (*p* p = 0.192, F = 1.8), and a significant effect of donor vs recipient (*p* < 0.001, F = 9.1) with no interaction (**Figure 2d**). Pairwise comparisons found that while fecal samples from mice receiving pre-shipping material from B6N had not reached full diversity at one week post-FMT, no differences were detected between either recipient group and their B6N donors in fecal or cecal diversity at two weeks post-FMT. Collectively, these data suggest that while the GM of B6N mice is more efficient at comprehensive colonization of GF mice than the GM of B6J mice, both GMs mature within recipients and develop alpha-diversity comparable to their donors. These data also suggest that the B6J GM is preferentially susceptible to the effects of shipping.

### Beta-diversity in recipients replicates effects of shipping on donor microbiome

Beta-diversity among the entire dataset was dominated by the differences between B6J and B6N donors and their respective recipients (**Supplemental Figure S2**). To better resolve the effects of shipping, data were stratified by donor substrain and analyzed separately. Within both donor substrain and their respective recipients, samples separated along PCo1 according to their status as donor or recipient (**Figure 3a, b**). Notably, within B6J and their recipients, examination of PCo3 revealed complete separation of recipient samples from pre-or post-shipping samples from the same mice at both timepoints and in cecal contents (**Figure 3c**). No such separation was observed among recipients of FMT from B6N donors (**Figure 3d**) on PCo3 or subsequent coordinates. Two-way PERMANOVA confirmed significant differences between donors and recipients in both donor substrains, but significant effects or interactions of donor shipping were only detected in B6J mice, again suggesting selective effects of shipping on the B6J donors and recipients. Lastly, rather than comparing groups, we determined the specific Jaccard distance between each recipient and their specific donor sample, and then assessed the effect of substrain, shipping, and timepoint post-FMT on mean donor-recipient distance using a three-way ANOVA. No significant effect of any factor was detected (substrain: *p* = 0.28, F = 1.2; shipping: *p* = 0.291, F = 1.2; timepoint: *p* = 0.661, F = 0.4).

**Figure 3.**
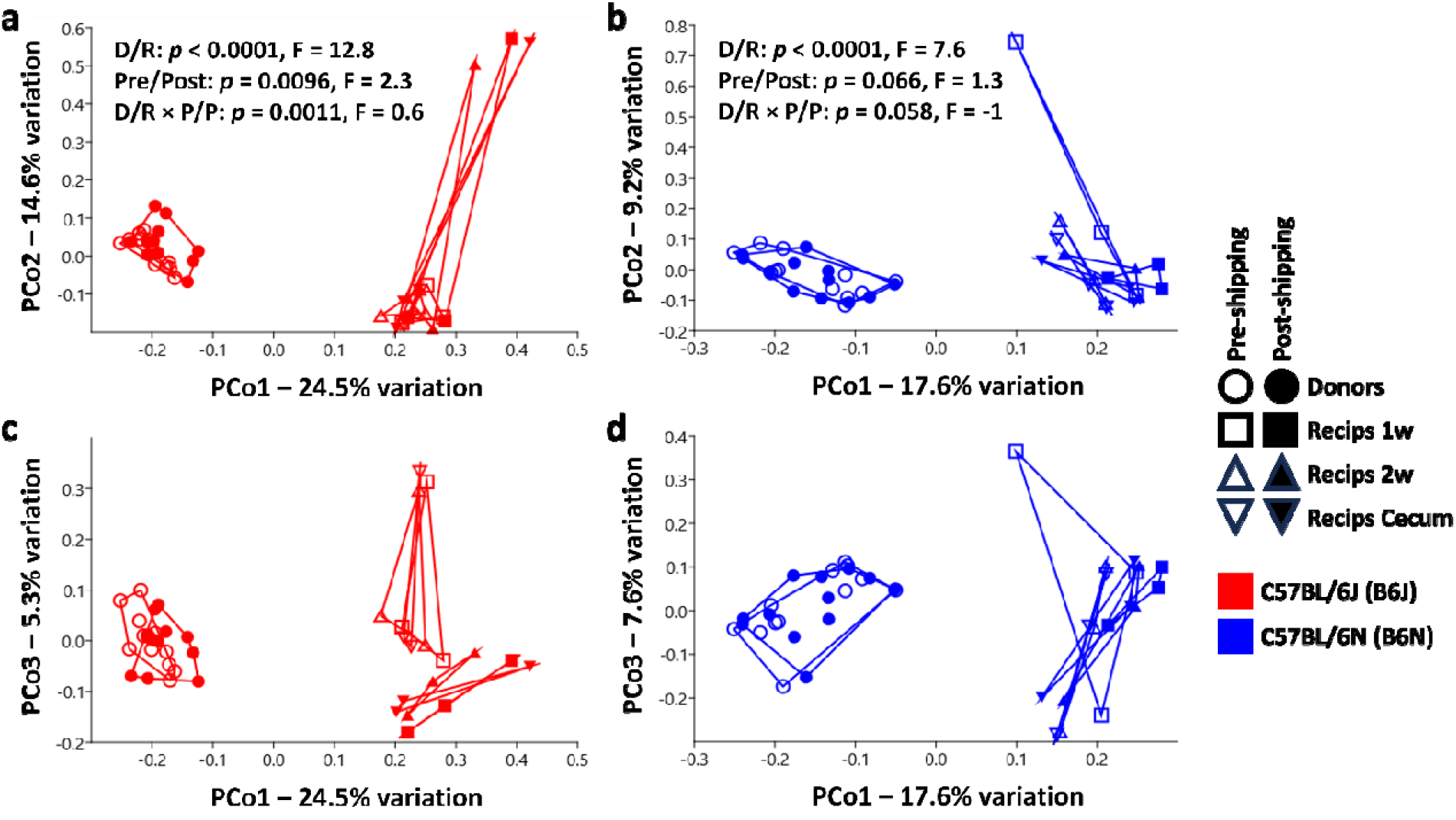
Principal coordinate analysis plots showing similarity between pre- and post-shipping C57BL/6J **(a)** or C57BL/6N **(b)** donor material and their respective recipient feces at one week (1w) or two weeks (2w) post-FMT or cecal contents. **(c, d)** the same data ordinated using PCo1 and PCo3, showing the separation of B6J recipients based on the timing of donor sample collection.

## Discussion

The current study was designed to determine whether the effects of shipping on the microbiome of donor mice resulted in similar changes in the microbiome of recipient mice receiving donor feces via fecal microbiome transfer (FMT). A secondary question was whether the potential effects of shipping on donor and recipient microbiomes were consistent across different SPF mouse microbiomes. Using C57BL/6 substrains purchased from the Jackson Laboratory or Envigo as donors, we used GF Swiss Webster mice as recipients, rather than antibiotic-treated pseudo-GF mice to eliminate any effect of differences in the recipient microbiome pre-FMT. As previously reported^10^, shipping resulted in subtle changes in the GM of donor mice. However, the current data also suggest that the effect of shipping on donor and recipient microbiomes is not consistent in both SPF microbiomes. A greater proportion of the Envigo GM successfully colonized the recipient gut compared to the Jackson GM, and the effect of shipping donor mice was less apparent in recipients of FMT from Envigo donors. These outcomes are counterintuitive given the substantially greater richness of the Envigo GM.

Despite those findings, the mean Jaccard distance between donors and recipients did not differ between substrains or use of pre-versus post-shipping samples. Thus, the differences observed between recipients of B6J fecal material collected from pre- and post-shipping donors, likely reflect the effects of shipping on the donor microbiome, as opposed to true differences in transfer efficiency.

The major limitation of this work is the limited sample size, which was directly related to the cost of GF recipient mice. Attempts were made to maximize the value of GF recipients through sampling at multiple timepoints and use of two SPF microbiomes. The low sample size is of greatest concern in the interpretation of negative findings, and the significant differences detected between groups are compelling based on the similarity of outcomes in B6J and B6N, or across multiple recipient timepoints.

In a real world scenario wherein banked fecal samples are needed in order to characterize the historical microbiome of a mouse colony, or colonize a different group of mice, these data indicate that donor samples may be subtly influenced by shipping in some mice, but that the effect of shipping is minor compared to other biological differences such as supplier-origin (SO) GMs. As mentioned earlier, only a handful of genetic backgrounds are readily available as GF mice, and scenarios in which the GM needs to be restored in a live colony would likely require antibiotic-mediated depletion of the endogenous GM prior to FMT. Previous work has demonstrated the hierarchical nature of SO GMs, exemplified by an inability to deplete and supplant the Envigo GM with the less rich Jackson GM, and faithful, complete transfer in the reciprocal direction^22^. Thus, effective reconstitution of live colonies may require rederivation or cross-foster approaches in some cases.

## Funding acknowledgement

This work was supported by U42 OD010918 from the NIH, Office of the Director to the Mutant Mouse Resource and Research Center at the University of Missouri (MU MMRRC).

## Statement of data availability

All sequencing data supporting the current research project are available in the National Center for Biotechnology Information (NCBI) Sequence Read Archive (SRA) as BioProject PRJNA1083537.

---

**Supplemental Figure S1.**
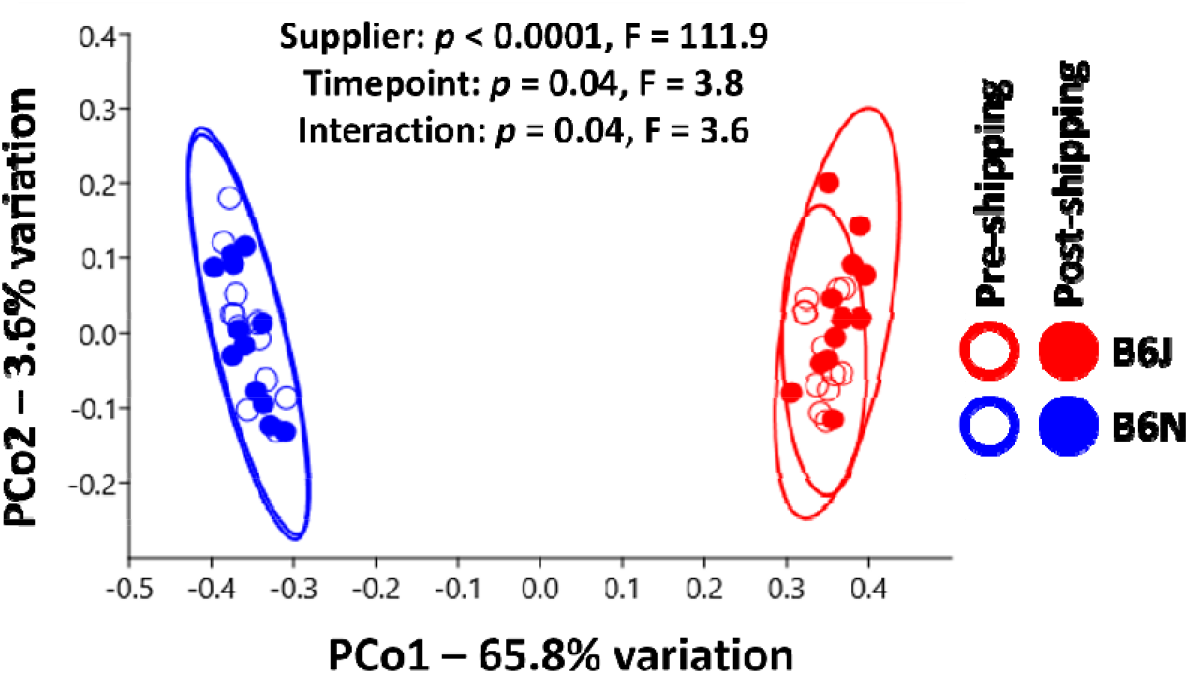
Principal coordinate analysis plot based on Bray-Curtis distances, showing separation of B6J and B6N donors, and overlap within each substrain of samples collected pre- and post-shipping; legend at right. Results of two-way PERMANOVA using Bray-Curtis distances.

**Supplemental Figure S2.**
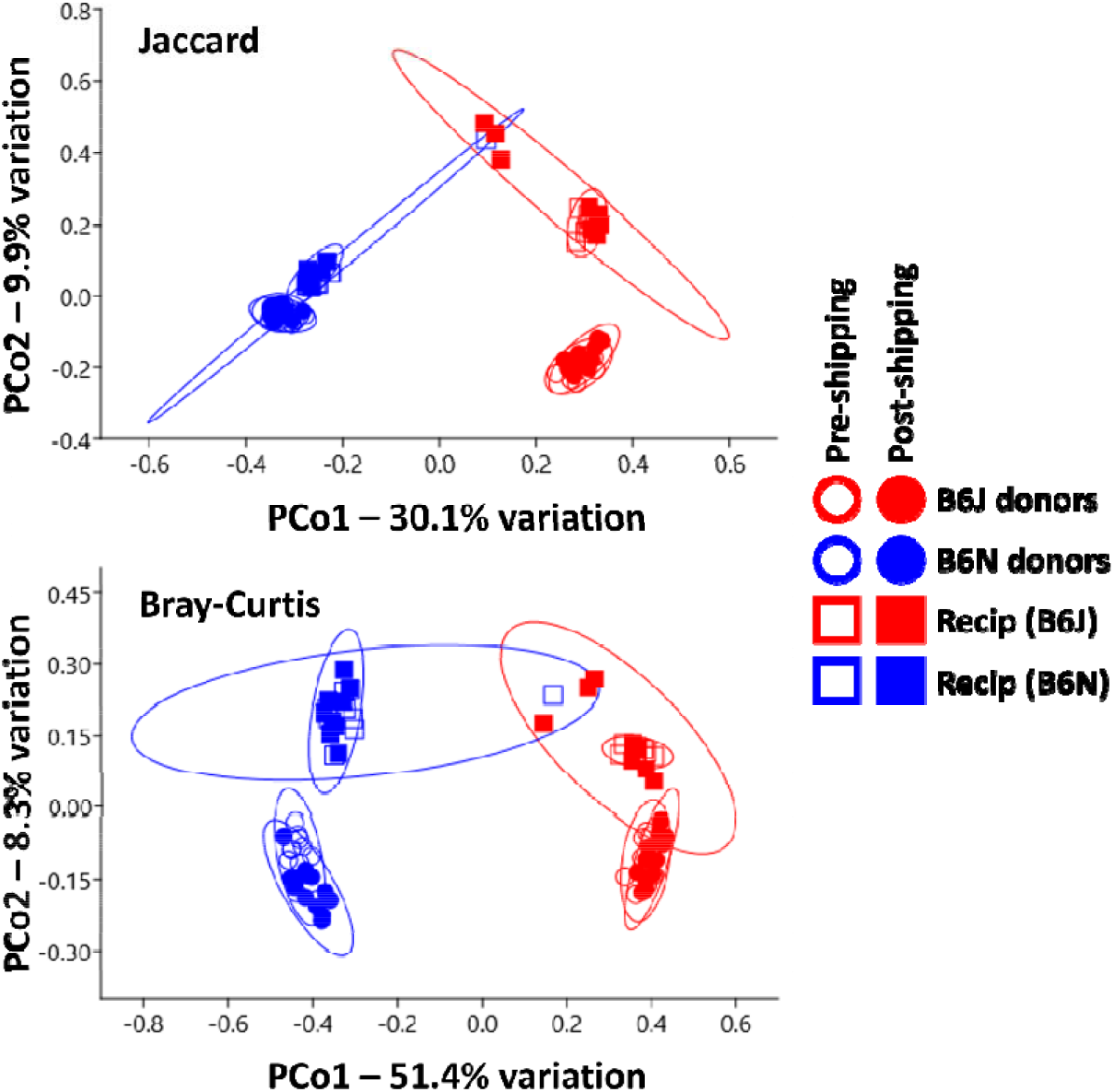
Principal coordinate analysis plots based on Jaccard **(a)** or Bray-Curtis **(b**) distances, showing the primary separation of C57BL/6 substrain (B6J, B6N) donors and their respective recipients along PCo1, secondary separation of donors and recipients along PCo2.

